# Comparing metabolic engineering scenarios using simulated design-build-test-learn-cycles

**DOI:** 10.64898/2026.02.03.703462

**Authors:** Paul van Lent, Sara Moreno Paz, Joep Schmitz, Thomas Abeel

## Abstract

Design-Build-Test-Learn (DBTL) cycles are a widely employed engineering framework in metabolic engineering. Nonetheless, their performance depends on a wide range of experimental and algorithmic design choices, whose combined effects on the successful optimization of microbial strains remain an open question. In this study, we performed *in-silico* DBTL cycles based on metabolic kinetic models to quantitatively assess how key process parameters affect strain optimization outcomes across four distinct metabolic pathway models. This includes parameters governing DNA library design, experimental budget limitations, and machine learning configuration. The results show that screening capacity is a dominant driver of optimization success, whereas DNA sequencing capacity has surprisingly little impact, despite its importance for model training. Selecting top-producing strains for sequencing consistently outperforms stratified sampling, highlighting a trade-off between predictive accuracy and optimization efficiency. DNA library structure strongly affects performance: increasing the number of editable positions generally improves outcomes, while expanding the set of gene targets can hinder optimization due to increased dimensionality or sparse sampling. Together, these findings offer actionable guidance for designing more effective DBTL workflows and underscore the value of simulation frameworks for exploring metabolic engineering strategies prior to experimental implementation.

## 1 INTRODUCTION

Metabolic engineering aims to improve microorganisms by genetically modifying their metabolic pathways to boost the synthesis of desired products Cho et al. (2022). This can be used to develop microbial cell factories that facilitate the synthesis of bulk and specialty chemicals using potentially sustainable substrates Tickner et al. (2021). However, this development from a proof-of-principle strain to a microbial cell factory is a complicated process, often taking many years of development and significant financial investment Beall et al. (2019). Reducing strain development times is essential for the widespread adoption of bioprocesses for manufacturing.

Advances in high-throughput screening and DNA sequencing technologies have allowed for the targeting of multiple metabolic pathway elements simultaneously for optimization Dietrich et al. (2010). This approach is known as combinatorial pathway optimization Jeschek et al. (2017). By targeting many elements in parallel, the likelihood of finding an optimal configuration for the production of a desired compound increases compared to the sequential de-bottlenecking of metabolic pathways. However, as the number of pathway elements targeted increases, the number of theoretical designs combinatorially explodes. To navigate this large landscape of possible designs, the Design-Build-Test-Learn (DBTL) cycle is widely used in metabolic engineering. In the Design phase, a set of designs is chosen based on the selection of targets and experimental design. This is then followed by the Build and Test phase, during which the strains are constructed and screened for their performance. The results from the screening phase are used to guide new designs in the subsequent DBTL cycle (Learn phase). This iterative approach has been shown to be successful in many applications Opgenorth et al. (2019); Hägele et al. (2025). However, due to the flexibility of the DBTL cycle framework, many open questions remain regarding how DBTL cycle process parameters affect strain optimization performance.

Machine learning (ML) has been increasingly used to help propose new strain designs for subsequent DBTL cycles Radivojević et al. (2020); Yang et al. (2025); Wei et al. (2025). These ML-assisted approaches are promising for automated strain engineering, as they effectively close the loop between DBTL cycles in a data-driven manner HamediRad et al. (2019). The success of the recommendation strategy used is closely linked to the process parameters related to the DBTL cycle, such as the sequencing and screening budgets, the design choices of the DNA library Jeschek et al. (2016), and the optimization exploration-exploitation tradeoffs. Often, the parameters of the DBTL cycle process are a consequence of experimental budgets, lab capabilities, and design choices that stem from experience. Consequently, elucidating how ML-assisted optimization performance depends on the experimental design of the DBTL cycle is critically important for the successful implementation of strain engineering.

*In silico* experiments serve as a versatile tool for evaluating metabolic engineering scenarios that would not be practically feasible in an experimental setting Moreno-Paz et al. (2024a); van Lent et al. (2023); Roy et al. (2021). Previously, we developed simulated DBTL cycles, a framework based on kinetic models to consistently evaluate ML methods across different metabolic engineering scenarios van Lent et al. (2023). This framework provided insights into ML methods, data requirements, and budgetary constraints, although it was limited to one metabolic model and only a few DBTL cycles. Whereas numerous studies have examined the machine-learning components of recommendation strategies Wei et al. (2025); Sabzevari et al. (2022); HamediRad et al. (2019), this work instead focuses on how the process parameters of the DBTL cycle influence the optimization performance of an ML-assisted strategy. Four bioprocess models with hypothetical metabolic pathways of diverse topologies and complexities are constructed and used as a test suite. The goal is to address experimental considerations and their influence on strain optimization, demonstrating their utility in refining experimental design for industrial metabolic engineering. We offer code and workflows for user-specific simulated DBTL cycle scenarios at https://github.com/PHvanLent/BenchmarkProjectDSMF/tree/main/scripts.

## 2 MATERIALS AND METHODS

### 2.1 Models

#### 2.1.1 General information

Four bioprocess models of varying complexity and scale were constructed using the *jaxkineticmodel* Python package van Lent et al. (2025a) and subsequently exported to SBML format Bornstein et al. (2008) for standardized representation and downstream analysis. Each model describes a hypothetical metabolic pathway designed to satisfy carbon balance across all reactions. Biomass formation reactions were parameterized to yield physiologically reasonable biomass yields of approximately 0.5 gram biomass per gram glucoseVan Hoek et al. (1998).

To account for the metabolic burden associated with genetic modifications, a protein load term was incorporated Wu et al. (2016). Metabolic burden was represented by an increase in the maximum rate of a non-growth-associated maintenance reaction that depletes ATP required for biomass synthesis. Specifically, the parameter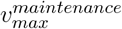 in the maintenance ATP sink reaction (eq. 1) was scaled relative to a wild-type reference strain. In the reference condition, the maximum flux is given by 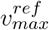, where all pathway enzyme concentrations [*E*_*i*_] are normalized to one. A constant *C* was introduced to ensure that the resulting fraction evaluates to unity under the reference enzyme concentration profile, thereby preserving 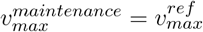 for the wild-type model while enabling systematic variation of metabolic burden in perturbed strains.

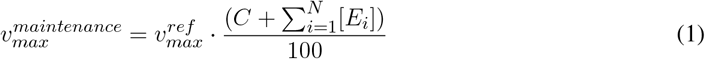

#### 2.1.2 Model statistics

Table 1 summarizes the structural properties of all pathway models, including the numbers of species, reactions, parameters, and optimization targets. To quantify metabolic complexity, we employed the method described in Ghavasieh and De Domenico (2024). Briefly, a graph Laplacian is constructed from the stoichiometric matrix *S* (eq. 2), which defines a random walk of duration *τ* over the reaction network. The *n*-th step of this random walk yields a matrix *M*, from which we compute the density matrix *ρ*_*τ*_ by normalization with respect to the trace of *M* (eq. 4). The von Neumann entropy of *ρ*_*τ*_ (eq. 5) is then used as a scalar measure of network complexity. The walk length was set to *τ* = 10, which we verified to be sufficiently long for the random walk entropy to reach a steady state. This procedure provides a consistent metric for comparing the complexity of distinct metabolic networks (see SI C). To enable comparisons across a diverse set of metabolic pathways, we constructed multiple topological variants for each pathway. Detailed model formulations, parameterizations, and validation procedures for all four models are provided in SI A.

**Table 1.** Key statistics of the four pathway models used in this study. The network complexity is a random walk measure for the stoichiometric matrix *S*. The topology characteristic gives a brief description of the features in the model.

| Model | Species | Reactions | Parameters | Enzyme targets | Network complexity | Topology characteristic |
| --- | --- | --- | --- | --- | --- | --- |
| Pathway A | 23 | 23 | 86 | 20 | 0.30 | linear |
| Pathway B | 18 | 16 | 59 | 12 | 0.32 | cycle |
| Pathway C | 25 | 21 | 82 | 17 | 0.22 | cycle + linear |
| Pathway D | 29 | 25 | 98 | 21 | 0.61 | cycle + linear |

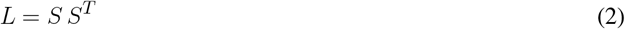

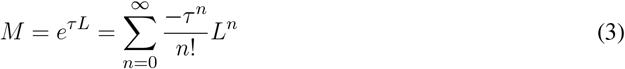

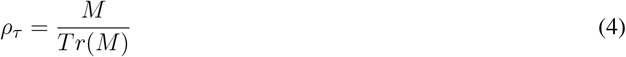

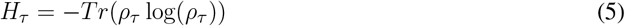

### 2.2 Metabolic engineering process

#### 2.2.1 Simulated Design-Build-Test-Learn cycles

The simulation of DBTL cycles was introduced in van Lent et al. (2023) and subsequently re-implemented using JAX Bradbury et al. (2018); van Lent et al. (2025a). The number of enzyme targets (*F* ) and promoters (*X*) for a set of library positions (*P* ) is used to construct a DNA library. This DNA library is represented as a matrix with *F*×*X* rows by *P* columns, where each element corresponds to a promoter-gene pair. A probability is assigned to each element such that the positions sum to one. Strains are constructed based on random assembly, according to these probabilities, so that a single strain includes *P* promoter-gene pairs. This approach mirrors the method used in many combinatorial pathway optimization methods involving library transformation Moreno-Paz et al. (2024b); Jeschek et al. (2017). In the first DBTL cycle, all promoter-gene pairs are equiprobable. In subsequent cycles, probabilities are adjusted using an ML-assisted approach (see next section). For all strains that are constructed (*S*) during the build phase, the designs are numerically simulated using Diffrax with the Kvaerno5 numerical solver Kidger (2022); Kværnø (2004). In the test phase, the built strains are simulated as a batch bioprocess without additional glucose feeding. Then, 10% homoscedastic noise is introduced. Finally, these simulated strains serve as training data for a ML model, which will be used in the next DBTL cycle.

#### 2.2.2 Recommending new strains: an ML-assisted approach

To automate the recommendation of new strains for comparing metabolic engineering scenarios across multiple DBTL cycles, we use an ML-assisted recommendation method that outputs a probability distribution for the DNA library that will be used in the next DBTL cycle.

##### 2.2.2.1 Machine learning model

XGBoost is trained on the input strain design data and the simulated values of the product of interest Chen et al. (2015). Default hyperparameters were used, with the exception of number of boosting rounds and earlystopping conditions (*num boosting rounds*=20, *early stopping rounds*=40). 10-fold cross-validation was performed on 20% of the training data, and the Pearson correlation coefficient was used as a performance estimate.

##### 2.2.2.2 Recommendation

The DNA library was randomly sampled during each DBTL cycle to generate a large candidate set (600,000 variants). This down-sampling is necessary, as the combinatorial design space can be prohibitively large. Samples were then evaluated using the trained XGBoost model. The predicted strain performances were expressed relative to the parent strain and subsequently converted into a probability distribution using the softmax function (eq. 6) Sutton et al. (1998). In this formulation, the parameter *β* controls the trade-off between exploration of the DNA library design space and exploitation of the model predictions.

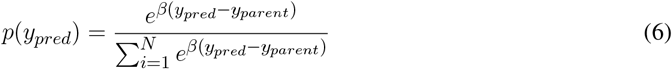

Strains predicted to outperform the parent strain are assigned higher probabilities than those predicted to underperform. To obtain probabilities for each DNA element in the library, we summed the probabilities from equation 6 over all strains containing that element. This procedure yields element-wise (column-wise) probability distributions across the DNA library, as determined by the ML model.

##### 2.2.2.3 Dynamic balancing exploration-exploitation

To automatically balance exploration and exploitation, we selected the softmax temperature parameter *β* using an entropy-based heuristic. For a given *β*, we computed the entropy of the softmax distribution (Equation 6) according to

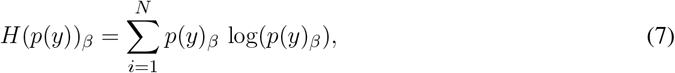

where *p*(*y*)_*β*_ denotes the softmax probabilities over the *N* candidates at temperature *β*. We evaluated *H*(*p*(*y*))_*β*_ over a range of temperatures (*β∈* [0, 100]) and identified the value of *β* at which d*H/*d*β* = 0. This point corresponds to the steepest decrease in entropy, which can be interpreted as the onset of a transition from a high-entropy (more exploratory) to a low-entropy (more exploitative) regime. The *β* value at this point was then used to compute the updated DNA library sampling probabilities in each iteration. This procedure was inspired by the need to avoid premature convergence to local optima in Bayesian optimization and reinforcement learning settings Sutton et al. (1998); Berger-Tal et al. (2014).

### 2.3 Metabolic engineering scenarios

To understand how the success of the DBTL cycle is impacted by design choices, the key DBTL cycle process parameters that were identified are reported in Table 2. Typical ranges for these parameters are shown, but they may vary depending on the specific pathway model (see code availability). For each model, we rank the enzyme importance using sensitivity analysis on the kinetic modelZi (2011). This serves as the ground truth ranking. Ten percent homoscedastic noise is added to the simulated production values.

**Table 2.** Description of process parameters.

| Symbol | Description | Model A | Model B | Model C | Model D |
| --- | --- | --- | --- | --- | --- |
| $F$ | Number of enzyme targets | 6-19 | 6-12 | 6-17 | 5-21 |
| $X$ | Number of distinct promoters | 2-6 | 4 | 4 | 4 |
| $P$ | Number of positions in library | 4-10 | 4-10 | 4-10 | 4-10 |
| $S$ | Number of screened strains | 50-1200 | 50-1200 | 50-1200 | 50-1200 |
| $N$ | Number of DNA sequenced strains | 50-200 | 50-200 | 50-200 | 50-200 |
| Sampling approach | DNA sequencing selection method | stratified or best sampling | stratified or best sampling | stratified or best sampling | stratified or best sampling |
| $\beta$ | Exploration-exploitation parameter | 0-100 | 0-100 | 0-100 | 0-100 |

Two rounds of computational experiments were conducted. In the first round, each individual process variable listed in Table 2 was evaluated for pathway model A to identify those that significantly influenced the optimization process. This is performed on one pathway model to reduce the computational load for more elaborate follow-up experiments. In the second round, informed by these results, a large combinatorial experiment was designed using the subset of variables deemed most likely to affect metabolic flux optimization performance. The variables included the number of enzyme targets (*F* ), the number of screened strains (*S*), the number of sequenced strains (*N* ), and the number of engineerable positions in the DNA library (*P* ).

For each pathway model, all feasible combinations of these variables were generated, resulting in between 144 and 320 distinct process configurations, depending on the model. For each configuration, five simulated DBTL cycles were executed. Performance was quantified by computing the area under the curve (AUC) of the production of the best built strain over five DBTL cycles and (ii) the median production across all built strains within each cycle. Each configuration was repeated for 10 independent simulation runs to account for stochastic variability. For certain pathway models, specific variable combinations could not be simulated (e.g., due to numerical stiffness or runtime errors); these configurations were excluded from subsequent analysis.

### 2.4 Code availability

Code and results are available on GitHub at https://github.com/PHvanLent/BenchmarkProjectDSMF.

## 3 RESULTS

### 3.1 Quantitative comparison of optimization processes reveals key DBTL cycle process parameters

Design-Build-Test-Learn (DBTL) cycles are widely used in metabolic engineering, helping to manage the extensive combinatorial design options of potential strain designs Jeschek et al. (2017). Despite its utility, the DBTL cycle involves many parameters that could affect the optimization of microbial strains. These include factors related to the experimental budget, experimental design, and ML-related factors. Given the substantial time and resources necessary for metabolic engineering experiments, employing simulated DBTL cycles offers a cost-efficient alternative to real-world experimentation for assessing process parameters in various scenarios Roy et al. (2021). In this research, we employ a previously developed kinetic model-based framework to assess the impact of process parameters on four distinct metabolic pathway optimization tasks van Lent et al. (2023).

We first evaluated whether an ML-assisted recommendation strategy outperforms a random DNA library transformation baseline Wei et al. (2025). In the baseline, each cycle constructs a library of strains by random sampling, and the best-performing strain from this library is selected as the parent for subsequent DBTL iterations. In contrast, the ML-assisted strategy uses a gradient bandit algorithm to iteratively update strain-selection preferences (probabilities) based on predicted productivity van Lent et al. (2025b); Sutton et al. (1998). This increases the chance of finding a higher best-performing strain that can be used for follow-up DBTL cycles. As expected, the ML-assisted recommendation method surpasses the baseline in terms of average production and the maximum producer in the set of built strains (Fig. 1A,B). To consistently quantify performance across scenarios, the area under the curve is used to measure general optimization performance 1C).

**Figure 1.**
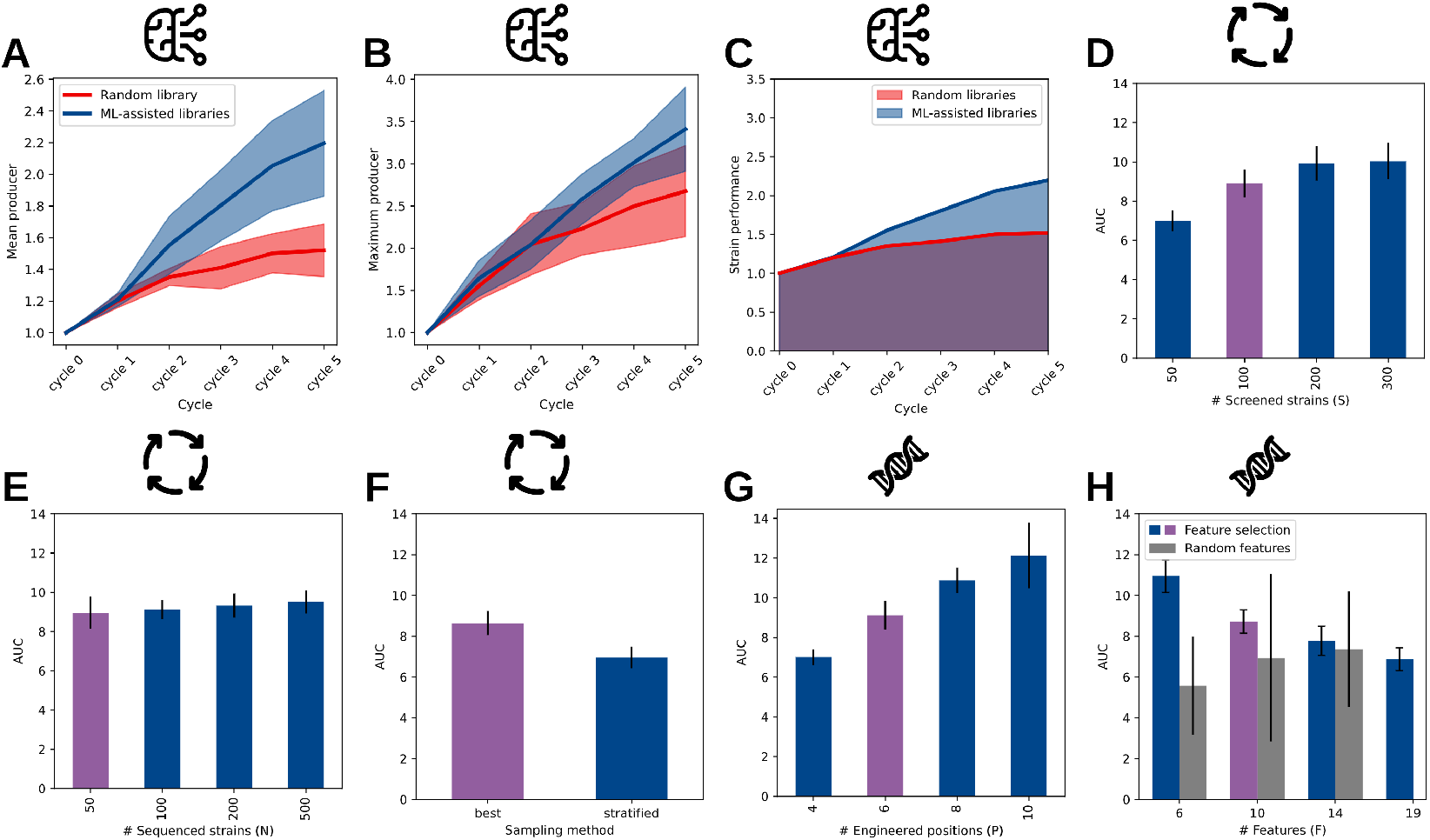
Evaluating process variables for pathway model. **A.** In plots D-H, the purple bar represents the baseline scenario (*S* = 100,*N* = 50,*F* = 10,*P* = 6,*X* = 4,*β* = 10). Three categories of process parameters are illustrated with pictograms: parameters related to ML, those related to the DBTL cycle, and parameters concerning library design. **A)** A comparison between a random library optimization approach and a strategy utilizing ML-assisted recommendations. The y-axis indicates the average production of the recommended strain set relative to the parent strain’s reference production. **B)** A similar graph, but depicting the top producer among the recommended strains. **C)** To assess scenarios, optimization performance is measured using the area under the curve (AUC). **D)** A comparison of approaches for selecting strains from screening results for DNA sequencing. The purple bar denotes the baseline scenario maintained across all plots. **E)** Optimization performance with varying DNA sequencing capacities. **F)** Sample selection strategies for screening: choosing the top producers (best sampling) versus a representative set (stratified sampling). **G)** The number of DNA library positions that can be integrated into the host strain’s genome.**H)** The impact of feature selection on optimization performance, with grey bars showing random feature selection.

Relating to DBTL cycle parameters, the impacts of screening capacity (*S*), DNA sequencing capacity (*N* ), and screening-to-sequencing strain selection are evaluated in terms of the area under the curve. Increasing screening capacity consistently improved optimization performance (Fig. 1D). This improvement likely occurs because larger screens increase the probability of including top-producing strains. Interestingly, increasing DNA sequencing capacity had little effect on optimization performance (Fig. 1E), despite the expectation that providing more data to the ML model would enhance predictive accuracy. We further compared two strain-selection strategies for sequencing: (i) selecting the top producers identified in screening (best sampling) and (ii) selecting a stratified set that spans the observed production range (stratified sampling). Best sampling led to notably better optimization performance compared to stratified sampling (Fig. 1F). In the case of stratified sampling, there is a chance that the best strain is not used as the new parent strain, which may result in worse optimization performance. Overall, these results suggest that expanding the screening capacity is more critical than DNA sequencing depth. Furthermore, choosing the best strains is preferred from a metabolic engineering optimization perspective.

Finally, we considered parameters associated with the DNA library, including the number of positions (*P* ) and the number of gene targets (*F* ). Findings suggest a significant effect of the DNA library positions on optimization performance by enhancing the exploration of the combinatorial design space (Fig. 1G). Although increasing the number of library positions might currently be technically challenging for strain engineering, these results underscore the potential benefits of developing synthetic biology tools that enable the construction and screening of larger libraries. Regarding the choice of the number of gene targets considered in the DNA library, reducing the number of targets significantly improves the optimization efficiency of ML-assisted recommendation (1H). However, this is only the case when the gene targets are chosen carefully, as selecting the wrong gene targets can result in worse performance than if no selection is performed (shown in gray). Consequently, feature selection methods, whether data-driven or grounded in mechanistic understanding, are crucial for improving metabolic engineering Van Rosmalen et al. (2024); Alonso-Gutierrez et al. (2015).

In summary, our observations indicate that optimization performance is significantly influenced by the screening capacity (*S*), the DNA sequencing selection technique, the number of library positions (*P* ), and the number of gene targets (*F* ). These findings will be further examined for a set of four unique metabolic pathway optimization problems in the sections below. Additional effects of other process parameters, such as promoter strength, exploration-exploitation trade-offs in ML-assisted recommendations, and the number of promoters included per gene, are reported in SI B.

### 3.2 Screening capacity, not sequencing capacity, is a key driver of optimization performance

Results found on pathway model A showed that increasing screening capacity greatly improves optimization, whereas increasing DNA sequencing capacity does not. To validate this finding, a grid search is performed for these two parameters.

Figure 2 presents four heatmaps for the different metabolic pathway models. The screening ratio indicates the down-sampling factor after screening. For instance, in a scenario involving 200 DNA-sequenced strains, a screening ratio of six leads to 1200 simulated strains. Across all models, an increase in the screening ratio correlates with enhanced AUC performance. Interestingly, the number of DNA-sequenced strains appears to have minimal effect on performance. While the overall performance differs between metabolic pathways, the independence of performance from DNA sequencing capabilities is generally observed. This is somewhat unexpected since the ML model is exclusively trained on this selection. It is often assumed that a larger training dataset would yield a superior model. In terms of optimization, the emphasis lies in identifying high-producing strains, which becomes more feasible with enhanced screening capabilities. In industrial metabolic engineering, the screening throughput of workflows may therefore represent a more significant bottleneck than DNA sequencing capacity.

**Figure 2.**
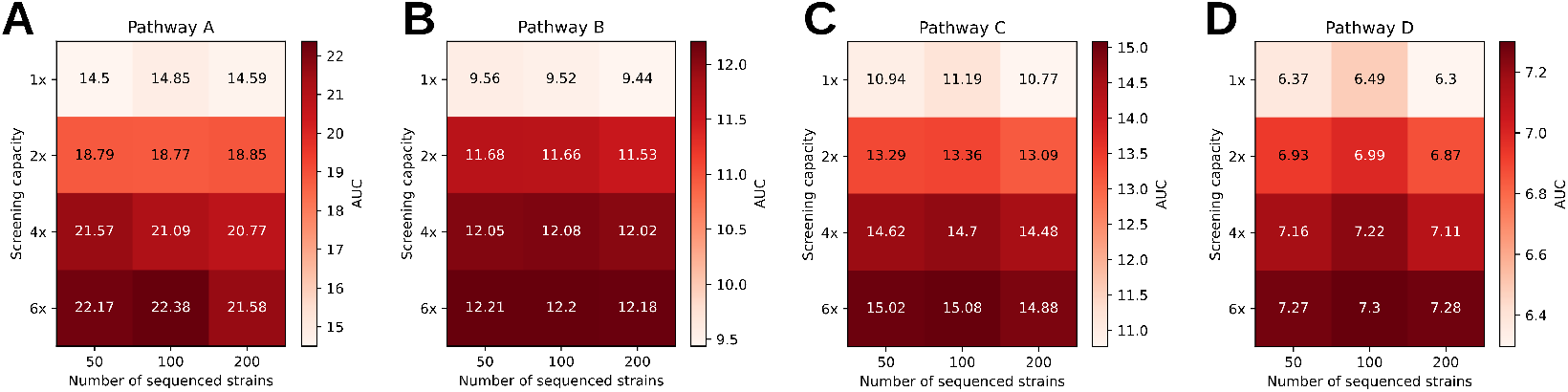
Screening capacity has a strong impact on ML-assisted optimization performance. **A)** Pathway model A (*F* = 6, *P* = 6). Screening capacity has a strong impact on the area under the curve performance, while DNA sequencing has little effect on the performance. **B-D)** A similar effect is observed for pathway model B,C,D.

### 3.3 Sampling top producers for screenings outperforms stratified approaches for metabolic flux optimization

Regression models in ML are susceptible to data imbalances stemming from selection biases, as noted in studies Yang et al. (2021); Tepeli et al. (2024); Steininger et al. (2021). In the field of metabolic engineering, these biases are often introduced when selecting which strains to sequence for further study. Ideally, from a metabolic flux optimization standpoint, one would select the highest-performing strains, even though they may not represent the underlying distribution well. To assess the implications of this selection, we compared the best sampling strategy (selecting top producers) with a stratified sampling method (see Materials and methods).

Across all pathway models, the selection of top producers consistently yields a higher AUC compared to using the stratified method (Fig. 3A-D). Notably, in metabolic pathway model D, after completing two cycles, the performance of the strain actually diminishes (Fig. 3D). This decline is attributable to the protein load effect accounted for in the kinetic models; as additional enzyme copies are included, biomass flux decreases due to increased maintenance. Nevertheless, selecting top producers in all DBTL cycles proves more effective than the stratified approach. Conversely, the performance of the ML model shows a slightly different trend. In pathway models A and B, the Pearson correlation coefficient on cross-validation sets is actually lower when using the best sampling method compared to the stratified method (Fig. 3E,F), which aligns with expectations for a biased distribution Steininger et al. (2021). In contrast, for pathway models C and D, no differences are observed between the two selection methods (Fig. 3G,H), potentially because the chosen strains more accurately reflect the underlying distribution.

**Figure 3.**
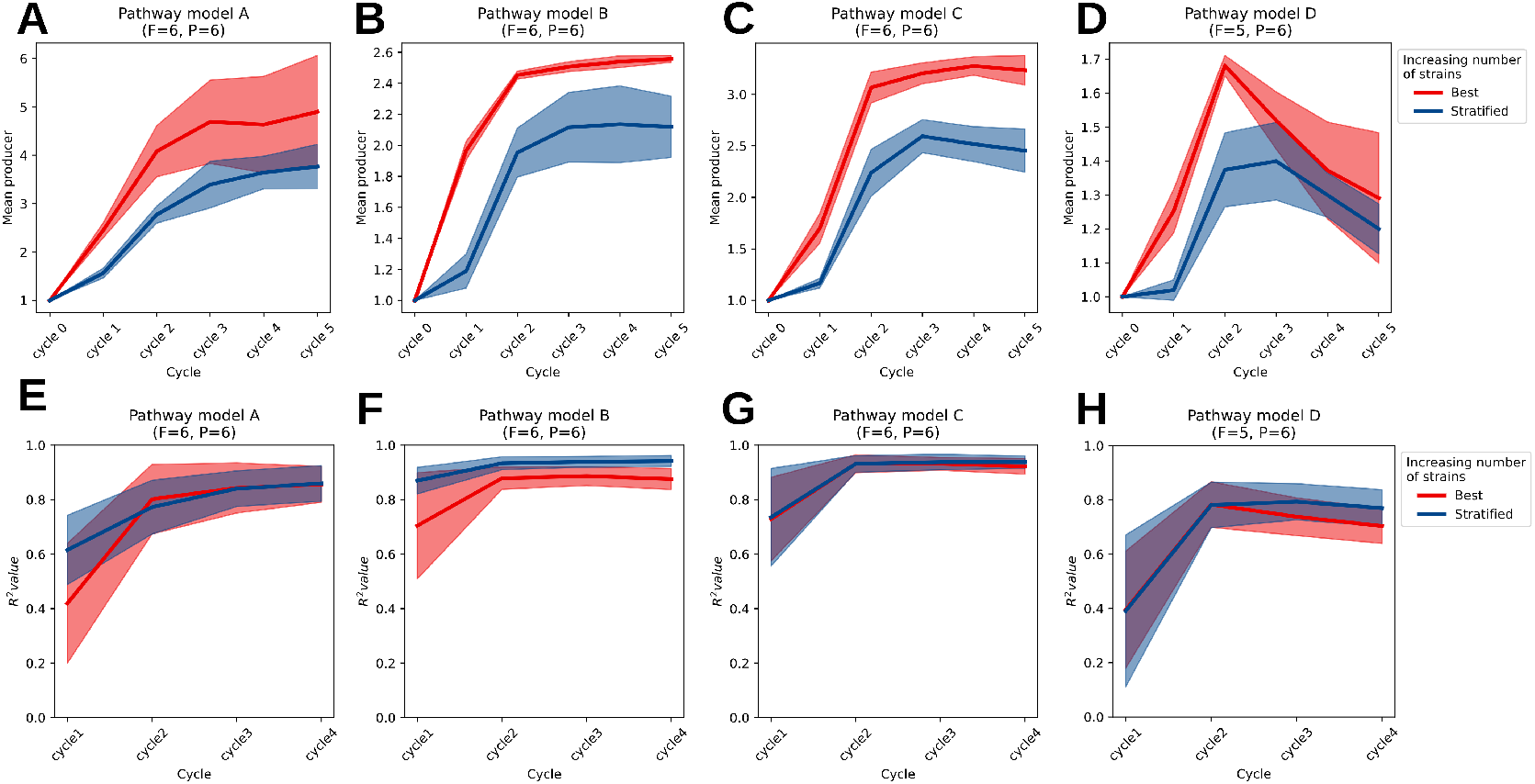
Selection method of strains for DNA sequencing. **A-D)** The optimization result for the pathway models. The stratified approach selects uniformly across the range of production values, while the best sampling approach selects the top producers (see Materials and methods). In general, product optimization favors a best sampling approach. **E-H)** The Pearson correlation coefficient *R*^2^ of the XGBoost model. A higher performance is typically observed for the stratified sampling approach.

To summarize, these results highlight a common trade-off in metabolic engineering. While a stratified selection approach improves the predictive accuracy of the XGBoost model, a bias towards selecting high-producing strains can result in better overall performance in strain optimization. Although we have chosen two distinct methods here, alternative selection strategies that consider both aspects may help to mitigate this trade-off.

### 3.4 DNA library parameters have mixed impact on strain optimization performance

In figure 1G and H, it was noted that parameters concerning the DNA library design have a considerable impact on performance. For a DNA library composed of a set number of positions (*P* ), each capable of accommodating different promoter-gene-terminator combinations (*X*×*F* ), the theoretical space of combinatorial designs is represented by (*F*×*X*)^*P*^ strains. Although enlarging the DNA library size can enhance genetic variation among strains, it also results in a combinatorial explosion of the design space. In this context, we aim to understand the effect of the number of included gene targets (*F* ) and the number of positions (*P* ) for the four pathway models.

We first evaluated how the number of gene targets in the library influenced performance across the four pathway models (Fig. 4A). Increasing the number of gene targets negatively affected performance for models A and C, whereas models B and D were mostly unaffected. This mixed effect might be explained for two reasons. First, as the dimensionality of the design space increases, the associated optimization problem becomes more challenging. Problems defined over larger gene sets are inherently more difficult to solve, as can be observed in models A and C Frazier (2018); Del Giudice (2021). Second, because each strain contains only six genes (*P* = 6), adding more candidate gene targets beyond this limit leads to sparser coverage of the combinatorial design space. This results in a less informative training dataset.

**Figure 4.**
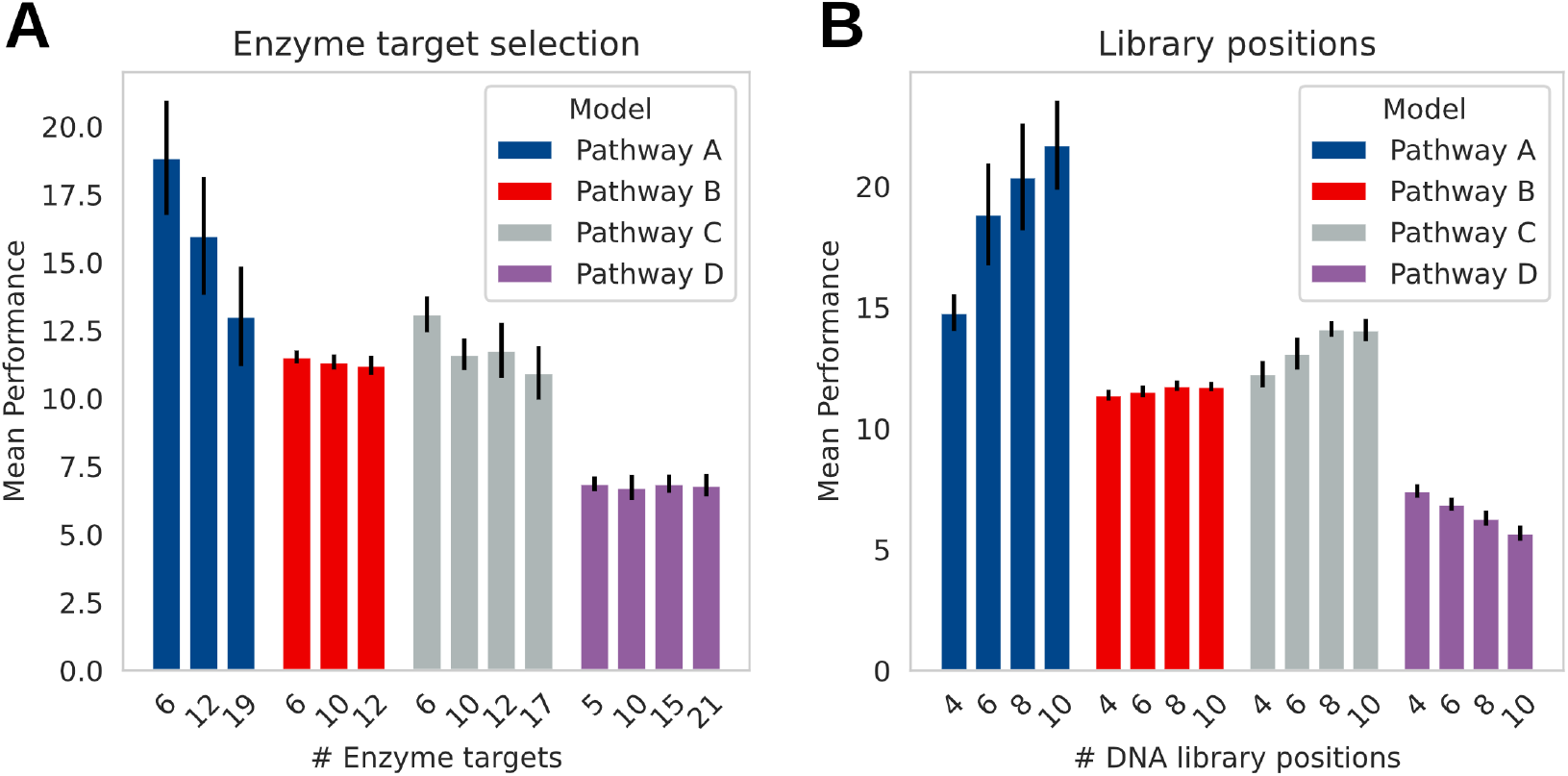
Library design: effect of feature selection and number of positions on optimization performance. **A)** Number of enzyme targets utilized for optimization in each pathway model. Enzyme targets were selected based on sensitivity analysis of the kinetic model Zi (2011). In general, fewer enzyme targets are associated with enhanced optimization performance. **B)** The number of DNA library positions that can be incorporated into a single strain. Typically, increasing the number of positions leads to improved performance, although it depends on the specific optimization issue (see Discussion).

The effect of increasing the number of library positions is shown in Figure 4B. For pathway models A and C, enhancing the number of library positions leads to a notable improvement in the AUC. Conversely, for pathway model B, such an increase does not result in any noticeable effect, indicating that only a limited number of genes may be crucial for performance enhancement. Interestingly, for pathway model D, the increased number of positions has a negative effect. This is likely due to the protein load effect included in the model. Introducing more gene copies reduces biomass production, thus hindering performance. This shows that while an expansion of engineering capabilities is potentially beneficial, it might depend on the specific pathway to be optimized.

The design of libraries is crucial in combinatorial pathway optimization Jeschek et al. (2017). Proposed strategies for DNA library design aim to balance genetic variation while preventing combinatorial explosion Jeschek et al. (2016). Here, we demonstrate that choosing appropriate genes and limiting the number of genes considered can enhance performance. Furthermore, the expansion of the number of DNA positions affects the area under the specific problem being addressed.

## 4 DISCUSSION

The DBTL cycle is a flexible and widely adopted framework for navigating the landscape of strain design options in metabolic flux optimization. In this study, we investigate how optimization performance depends on parameters associated with DNA library design and the DBTL cycle when using ML-assisted recommendation methods Moreno-Paz et al. (2024b); Pandi et al. (2022); Wei et al. (2025). To accomplish this, we employed simulated DBTL cycles, which offer a cost-effective alternative to real-world experimentation van Lent et al. (2023).

Our analysis began by identifying seven key parameters related to the DBTL cycle, specifically in the Design phase (number of features and positions), the Build and Test phase (screening capacity, sequencing capacity, and sampling approach), and the Learn phase (the exploration-exploitation trade-off) . These were tested on one pathway model to determine which parameters had a significant effect on optimization performance and should be further investigated (see Fig. 1 and Fig. **??**). This showed that screening capacity, the sampling approach for DNA sequencing, the number of positions in the library, and the number of features had the most prominent effect on performance.

An interesting finding when expanding the analysis to multiple pathways is that screening capacity, but not DNA sequencing capacity, was a major driver of optimization performance. While it would be expected that a model trained on more samples results in a better predictive model, this apparently does not translate into improved recommendations. We proposed that this occurs because more extensive screening raises the likelihood of identifying high-performing strains. As a result, the training data includes more observations that indicate regions of the combinatorial design space that may yield improved performance. This may also explain why sampling the best strains for DNA sequencing is a better strategy than the stratified approach, as the ML model observes more high-performing strains in this scenario. In this regard, expanding screening capacity can enhance the predictive performance of ML models, as it enlarges the domain and range of data on which the model is trained.

For gene target selection, enzyme sensitivities concerning the product were employed to rank and select the most important enzymes. In reality, performing gene target selection can be difficult, as it is not known *a priori* which genes affect production. Genome-scale modeling approaches could be used to choose gene targets Van Rosmalen et al. (2024); Domenzain et al. (2025). Expanding this analysis to other pathways indeed verifies that performance is positively impacted by feature selection, although the impact may differ between optimization targets.

For the number of library positions, we found that for pathway model D, the optimization performance decreases in contrast to the other models (Fig. 4B). This can be attributed to the network topology of pathway model D, which has a by-product route that might be favored over the product route (see SI A). Furthermore, it might also be a consequence of performing library transformation. As more copies of genes are added, the implemented protein load affects biomass formation, resulting in decreased production. This can also be observed in Figure 3D, where production drops after three DBTL cycles. Scenarios in which genes are down-regulated or knocked out as the main intervention could also be valuable to simulate Burgard et al. (2003), particularly for systems where metabolic burden poses a challenge. However,implementing down-regulation necessitates replacing native promoters in the host, which is more complex from a metabolic engineering standpoint. For this reason, we did not include it in the present study.

Balancing exploration and exploitation in machine-learning-driven recommendations within the DBTL cycle is known to be important Zhang et al. (2020); van Lent et al. (2025b); McInerney et al. (2018). In Figure 1, we fixed *β* to a single value to enable consistent comparisons between scenarios. In practice, the preferred exploration–exploitation balance may shift over the course of a combinatorial pathway optimization experiment. For instance, one might prioritize exploration in the early DBTL cycles and then gradually increase exploitation in later cycles. To avoid searching for this exploration-exploitation schedule, we applied an entropy-based approach to adaptively manage this trade-off (see Materials and methods). We observed that this strategy resulted in a minor improvement over a static approach. Nonetheless, alternative strategies for controlling exploration versus exploitation are conceivable and may be worth exploring in future work.

Overall, our results highlight practical considerations that can be used to improve the DBTL cycle for real-world metabolic engineering. As the variety of metabolic engineering scenarios that can be considered is abundant, workable templates for running alternative scenarios of the DBTL cycle are provided.

## AUTHOR CONTRIBUTIONS

## SUPPLEMENTAL DATA

### 4.0.1 SI A

This text contains plots of the validation and descriptions of the four models used in this study.

### 4.0.2 SI B

This text contains plots of process parameters from Table 2 that are not reported in the main results. These include the results of the exploration-exploitation parameter *β*, the number of promoters *X*.

### 4.0.3 SI C

This contains results on the network complexity of a set of metabolic networks.

## DATA AVAILABILITY STATEMENT

The datasets generated can be found in the 4tu repository (doi: **10.4121/f2833f74-2586-4632-8c44-7d98ead11a34**). Scripts and workflows can be found on the github page https://github.com/AbeelLab/BenchmarkProjectDSMF

