## Supplemental Figures for "Comparing metabolic engineering scenarios using simulated design-build-test-learn-cycles"

### *Supplementary Material for article Comparing metabolic engineering scenarios using simulated design-build-test-learn-cycles*

#### 0.1 Figures

##### 0.1.1 Model validation

##### 0.1.2 Additional plots for pathway model A DBTL cycle parameters

### 0.1.3 SI C

This contains results on the network complexity of a set of metabolic networks.

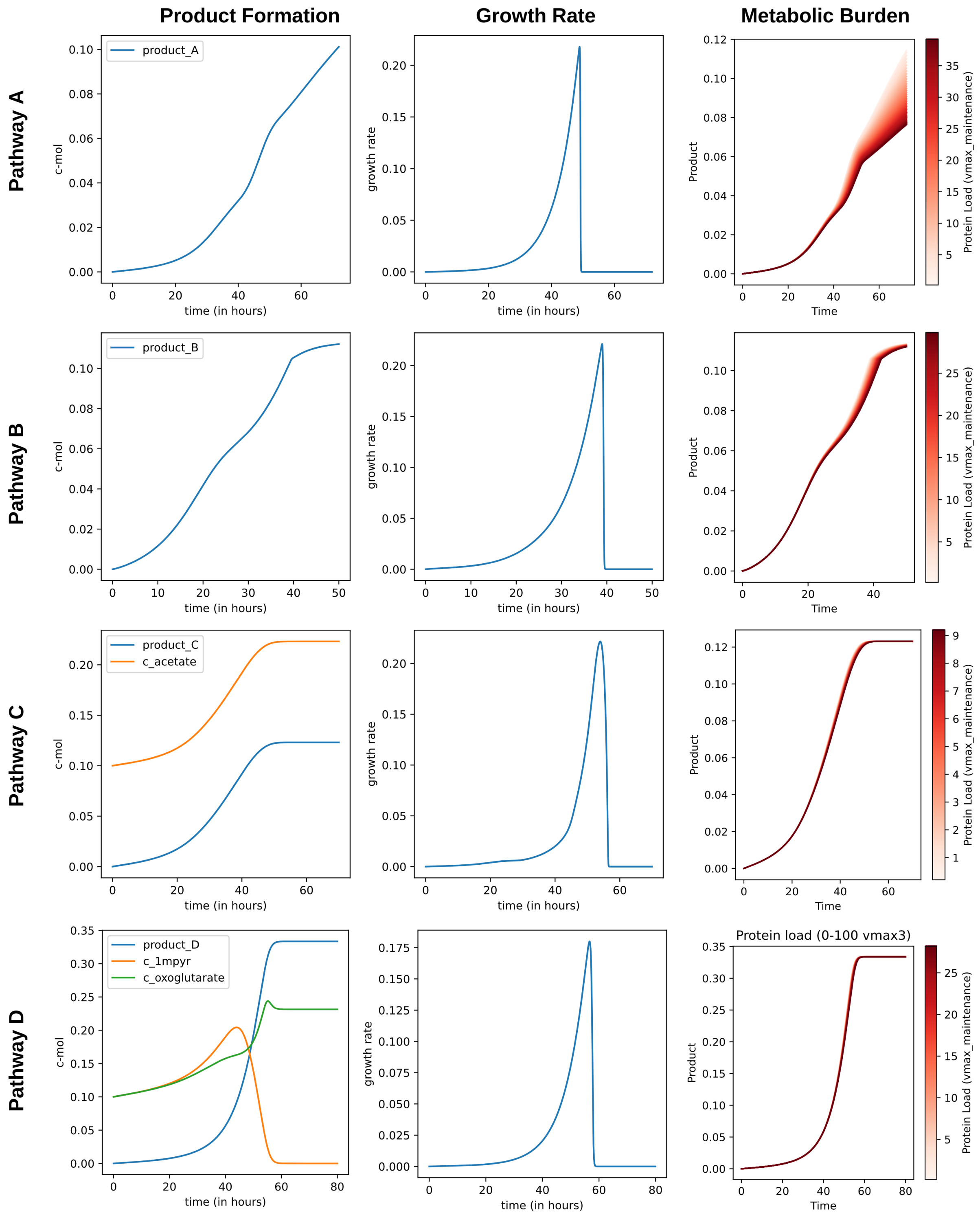


**Figure S1. Plots of some characteristics of the bioprocess models. Left)** Product formation in the four bioprocess models. Pathway C and D have byproducts. **Middle)** The growth rate for the four pathway models. **Right)** Effect of protein load on the four pathway models

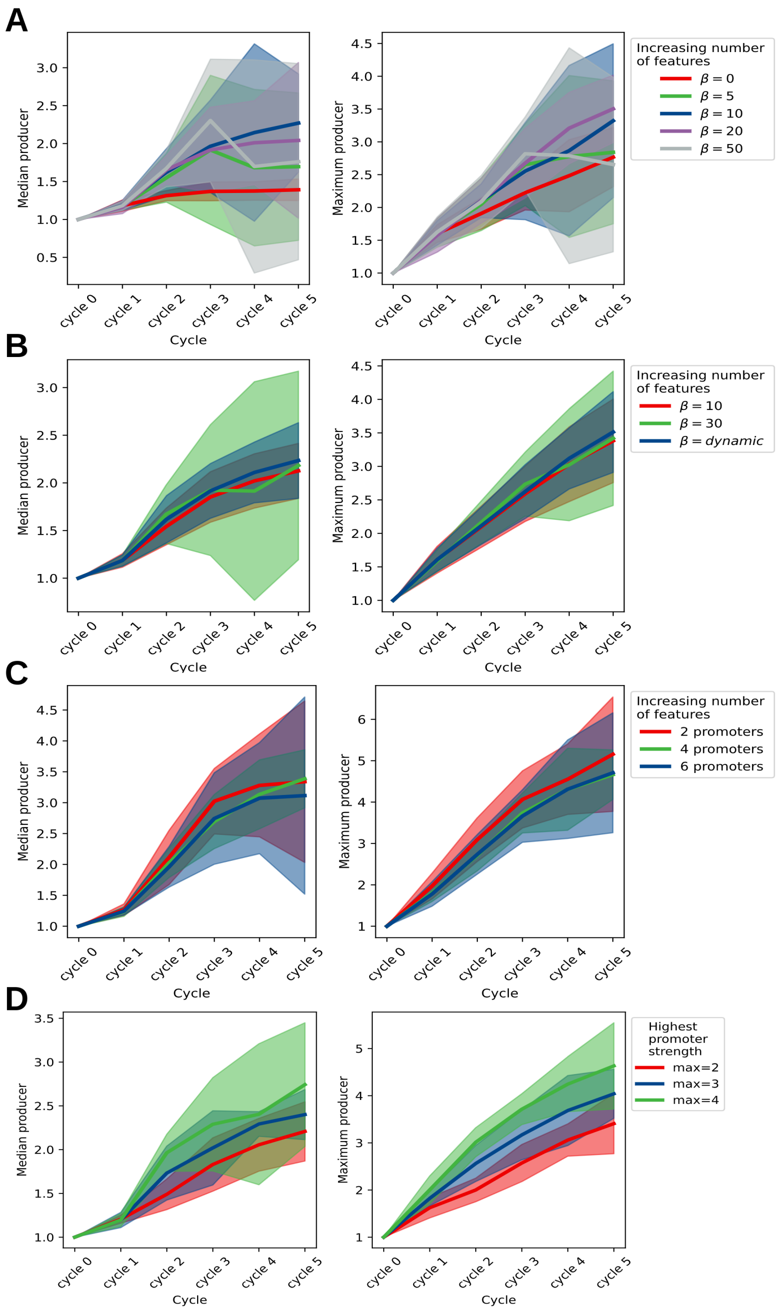


**Figure S2. Additional simulated scenarios for pathway model A**. **A)** The exploration-exploitation parameter $\beta$ and the effect on the median and maximum strain performance. **B)** Dynamically balancing this parameter does not improve optimization performance significantly. **C)** Number of promoters used in the library design, on the range 0-2. **D)** Scenario where the strongest expression strength is two, three, or four times stronger than the wild-type promoter.

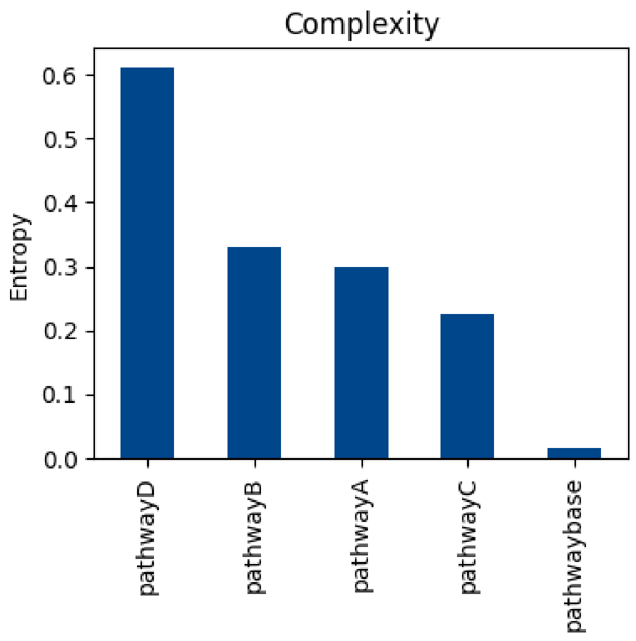


**Figure S3. Complexity of the metabolic pathway calculated using the approach taken in Ghavasieh and De Domenico (2024).** Pathway model D had the highest complexity associated, while pathway model C was considered the easiest in terms of topology.
